# Strong selection of a few dominant CD8 clones in a TLR7-dependent autoimmune mouse model

**DOI:** 10.1101/393819

**Authors:** Peter A. Morawski, Silvia Bolland

**Affiliations:** Laboratory of Immunogenetics, Division of Intramural Research, National Institute of Allergy and Infectious Diseases, NIH, Rockville, MD 20852, USA

## Abstract

Systemic lupus disease is characterized by the expansion of a self-reactive repertoire of B cells that, with the help of CD4 cells, generate IgG antibodies against common nuclear antigens. Meanwhile, the functional state and posible clonal selection of CD8 cells in lupus remain poorly defined. We previously described the activated but non-pathogenic phenotype of CD8^+^ T cells, some of which accumulate in the brain, in a model of systemic autoimmune disease triggered by increased copy number of the *tlr7* gene (TLR7tg mice). Here we report, through the analysis of TCRβ sequences, that CD8^+^ cells from TLR7tg are strongly selected for a small number of clones, some of them reaching 30% of the repertoire, compared to less than 0.4% for a single top clone in wild type cells. High frequency clones are variable in sequence among individual TLR7tg mice and are distinct from top clones in WT, while cells from spleen and brain-resident cells from the same animals have perfect concordance. These results suggest that top CD8 clones are selected in stochastic fashion in each animal but limit further diversification, and that brain infiltrating CD8 cells in TLR7tg are not selected by a tissue specific antigen. This kind of extreme clonal dominance and narrowing of the CD8^+^ T cell repertoire coud potentially impair anti-viral responses and should be considered as an additional detrimental feature of chronic autoimmune disease.

## Introduction

A substantial component of systemic autoimmune disease such as lupus is the breakdown of lymphocyte tolerance, primarily involving the expansion of autoreactive CD4^+^ T cells that provide help to B cells, which can in turn produce autoantibodies with various specificities (1). The contribution of CD8^+^ T cells to disease etiology or ongoing pathology is not well studied and no formal role for these cells has been established, though work from several groups now suggests CD8^+^ T cells could serve immunoregulatory roles in lupus (2).

We have studied lupus using several disease models including TLR7tg mice. These animals have a severe phenotype that includes glomerulonephritis, anemia, and a lymphoproliferative disorder that mainly impacts the CD4+ T cell and B cells pools, while the effect on the CD8 compartment is comparatively small (3, 4). We recently showed that the CD8^+^ T cells in these mice are non-pathogenic and have a strong effector phenotype compared to wild type controls, with the most activated cells accumulating in the brain (5). Because of the vast number of potential self-antigens available in both the periphery and brain, an understanding of the antigen-specific nature of these lymphocytes could help to better understand their relevance and function.

Membrane-bound antigen receptors are required by all lymphocytes for the recognition of peptide fragments bound to and presented by MHC proteins. The TCR serves as the specific antigen receptor on T lymphocytes and is composed of an alpha and a beta chain (6). The rearrangement of germline encoded variable (V), diversity (D), and joining (J) genes provides the immunological basis for the vast number of possible unique antigen receptors on lymphocytes. In naïve mice (7, 8) and in healthy humans (9, 10) certain common patterns of Vβ gene usage can be found in the formation of the TCR repertoire. Specific and narrow Vβ gene usage is evident in autoimmune disease models such as experimental autoimmune encephomyelitis (EAE) (11), and the systemic autoimmune models MRL/lpr (12) and B6-lpr (13), though these studies focus on CD4^+^ T cells and not CD8^+^ T cells. In lupus patients, an analysis of total T cell Vβ gene usage and the CDR3, which dictates antigen specificity, revealed significant changes in both compared to healthy controls, and revealed a narrowing of the overall TCR repertoire (14, 15). These data suggest that T cell antigen-specificity in lupus is important, yet we still understand little about the selective induction of autoreactive responses in this systemic disease.

In the current study, we address whether the peripheral and brain-resident CD8^+^ T cell populations in the TLR7tg model of lupus disease are selected by their antigen specificity. We were interested to understand whether these lupus-prone CD8^+^ T cells respond to environmental factors in a polyclonal fashion or if some strong selective pressures might exist that induce a more antigen-specific response. To accomplish this, we sequenced the TCRβ chains of this population of CD8^+^ T cells. Our results demonstrate strong clonal dominance of CD8^+^ T cells can arise in the context of systemic autoimmune pathology, suggesting that the function of these cells might be compromised in some severe cases of disease.

### Materials and Methods

## Mice and experimental protocols

The generation of TLR7tg mice has been described previously (4). All experiments in this study used the transgenic line 7.1 that harbors 8–16 fold TLR7 gene expression above wild type controls. All mice were maintained on a C57BL/6J background. Male mice between 12-13 weeks of age were used for these experiments. All animals were housed and studied in accordance with the approved NIH Animal Study Protocol, and all experimental protocols were approved by and performed according to NIH ACUC guidelines. All efforts were made to minimize animal suffering and to reduce the number of animals used.

### Vascular labeling of leukocytes

Discrimination of cells in the vasculature versus those in the brain parenchyma was performed as described (16, 17). Briefly, 5 μg of BV421-CD45.2 (1D4, BioLegend) diluted in sterile 1x PBS was injected i.v. once per mouse. Antibodies were allowed to circulate for a maximum of three minutes and then mice were euthanized and organs extracted. Subsequent *ex vivo* co-labeling of tissue-extracted lymphocytes with APCcy7-CD45.2 (1D4, BioLegend) is done as described below and allows discrimination between vessel (dual CD45.2 labeled) and parenchymal (*ex vivo* fluorophore only) cells.

### Lymphocyte extraction from tissue

Cell extraction from tissues was modified according to the described protocol (18). For lymphocyte extraction, organs were digested for 30 minutes at 37 °C, and shaken at 240 rpm in HBSS containing 1 mg/ml Collagenase Type I (Life Technologies). Digested slurry was pelleted, washed, and passed through a 70 μM mesh filter, then resuspended in Percoll (GE Healthcare) diluted to 90% using HBSS. Layers of 60% and 37% Percoll were then sequentially overlayed on the 90% cell-Percoll mix. Cell separation was accomplished by centrifugation at 8 °C, 500 × g for 18 minutes without brakes. Cells were washed and resuspended in staining buffer containing 5% FBS in HBBS in preparation for flow cytometry.

### Lymphocyte enrichment and flow cytometric sorting

Single-cell lymphocyte solutions were prepared from organs as indicated and resuspended in staining buffer containing 5% FBS in HBBS. Pre-enrichment of splenic lymphocytes was performed using PE-labeled CD8^+^ T cell magnetic bead positive selection according to the manufacturer instructions (RoboSep, Stem Cell). Splenic and brain-resident lymphocytes were identified using mAbs against the following antigens, all from BioLegend unless otherwise noted: PE-CD8α (MAR1), FITC-CD4 (RM4-5), BV711-CD11b (M1/70), BV421-CD45.2 (1D4), APCcy7-CD45.2 (1D4) APC-TCRβ, (eBio, H57-597). Samples were sorted on an AriaIII SORP (BD) equipped with violet (403 nm, 100 mW), blue (488 nm, 100 mW), yellow-green (561 nm, 50 mW) and red (638 nm, 150 mW) lasers, then analyzed with FlowJo (TreeStar Technologies). Pre- and post-sort purity of CD8^+^ T cells are shown in Figure 1A.

**FIGURE 1.**
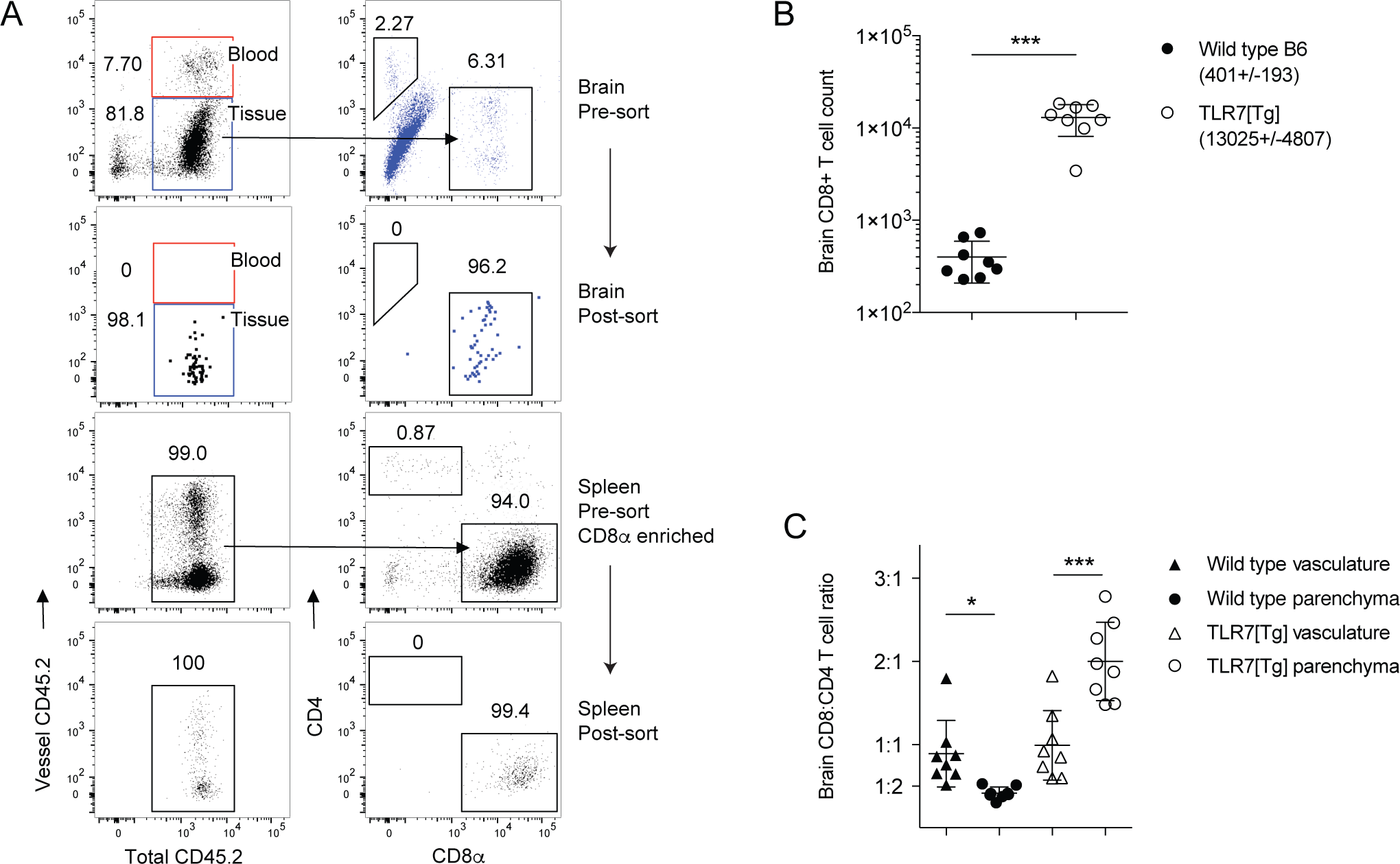
Isolating CD8+ T cells from lupus-prone mice. (**A**) Gating strategy (pre-sort) and results (post-sort) for flow cytometric isolation of CD8+ T cells from wild type B6 and TLR7[Tg] brain and spleen. Discrimination of cells in the parenchyma (tissue) from those in the vasculature (blood) was performed only for brain lymphocytes. Splenic lymphocytes were first positively pre-enriched for CD8α. (**B**) Absolute count of brain CD8+ T cells of animals from (A) with mean and s.d. listed. (**C**) CD8+/CD4+ T cell ratio in the brain of animals from (A). Error bars indicate mean + s.d. *P ≤ 0.05, **P ≤ 0.01, and ***P ≤ 0.001. Student’s t test, unpaired (B), paired (C).

### DNA extraction, TCRβ gene sequencing, and analysis

Frozen cell pellets of sorted wild type B6 and TLR7tg spleen (>2 x 10^5), and TLR7tg brain (>1 x 10^4) CD8^+^ T cells were sent on dry ice to Adaptive Biotechnologies (Seattle, WA) for genomic DNA extraction and subsequent TCRβ gene sequencing. Amplification and sequencing of TCRβ CDR3 was performed using the Adaptive Biotechnologies ImmunoSEQ Platform, which combines multiplex PCR with high throughput sequencing and a sophisticated bioinformatics pipeline for TCRβ CDR3 analysis (19, 20). Data analysis was performed using ImmunoSeq Analyzer 3.0 software also provided by Adaptive Biotechnologies.

### Statistical analysis

Statistical significance of data was calculated with Prism 6.0 software (GraphPad). For comparisons between two normally distributed groups a two-tailed unpaired *t*-test with Welch’s correction was used. For comparison between more than two groups statistical analysis was performed using a one-way ANOVA with the Tukey method. Gaussian distribution analysis of CDR3 sequences was used to characterize how population clonality as previously described (21). Error bars indicate mean + s.d. *P ≤ 0.05, **P ≤ 0.01, ***P ≤ 0.001, and ****P ≤ 0.0001 (Student’s t test).

## Results

### The CD8^+^ T cell repertoire in TLR7tg mice is oligoclonal

We previously described the presence of activated, although likely non-pathogenic, CD8^+^ T cells in the spleen and brain of TLR7tg that we hypothesized could regulate aspects of the systemic disease developed in these mice (5). To gain insight into the clonality of these cells, we performed TCRβ sequence analysis of CD8^+^ lymphocytes extracted from the spleens and brains of TLR7tg mice, or spleens of wild type controls. A combination of CD8-positive bead enrichment followed by flow cytometric sorting were used to select for CD8^+^TCRβ^+^CD4^−^CD11b^−^ T cells with a final purity of at least 99% (Fig. 1A).

Brain parenchymal CD8^+^ T cells were distinguished and separated from those in the vasculature using a combination of *in vivo* intravascular labeling and *ex vivo* staining as previously described (16, 17). A precise cell count provided during FACS isolation indicated an average of 13,025 CD8^+^ T cells per brain of each TLR7tg mouse compared with an average of only 401 in each wild type control (Fig. 1B). Because we required a minimum of 1,000 cells for effective TCRβ sequencing analysis, we did not to include the wild type brain CD8^+^ T cells in our study. Consistent with our previous findings (5), CD8^+^ T cells outnumbered CD4^+^ T cells in the brain parenchyma by 1.5- to 3-fold, a unique skewing not present in the vasculature or in other organs of lupus-prone mice (Fig. 1C). Genomic DNA from isolated CD8^+^ cells was used for TCRβ sequencing with the Adaptive Biotechnologies ImmunoSEQ Platform. The number of unique in-frame rearrangements per sample in TLR7tg spleens was decreased compared to wild type spleen (Fig. 2A) despite the total cell input (~2 x 10^5) and sum of all productive templates being the same (Fig. 2B). These data can be collectively summarized using a metric to measure total sample diversity, indicating that the peripheral TLR7tg CD8^+^ T cell repertoire has an increased degree of olgioclonality with respect to wild type controls (Fig. 2C). Similar oligoclonality was found in the brain-resident CD8^+^ T cell pool (Fig. 2A-2C) indicating that as a population these cells are largely the same as those in the periphery. Importantly, an equivalent percentage of cells sequenced from each of the three groups provided detectable productive rearrangements indicating equivalent quality of input DNA, sample handling, and sequencing efficiency (Fig. 2D).

**FIGURE 2.**
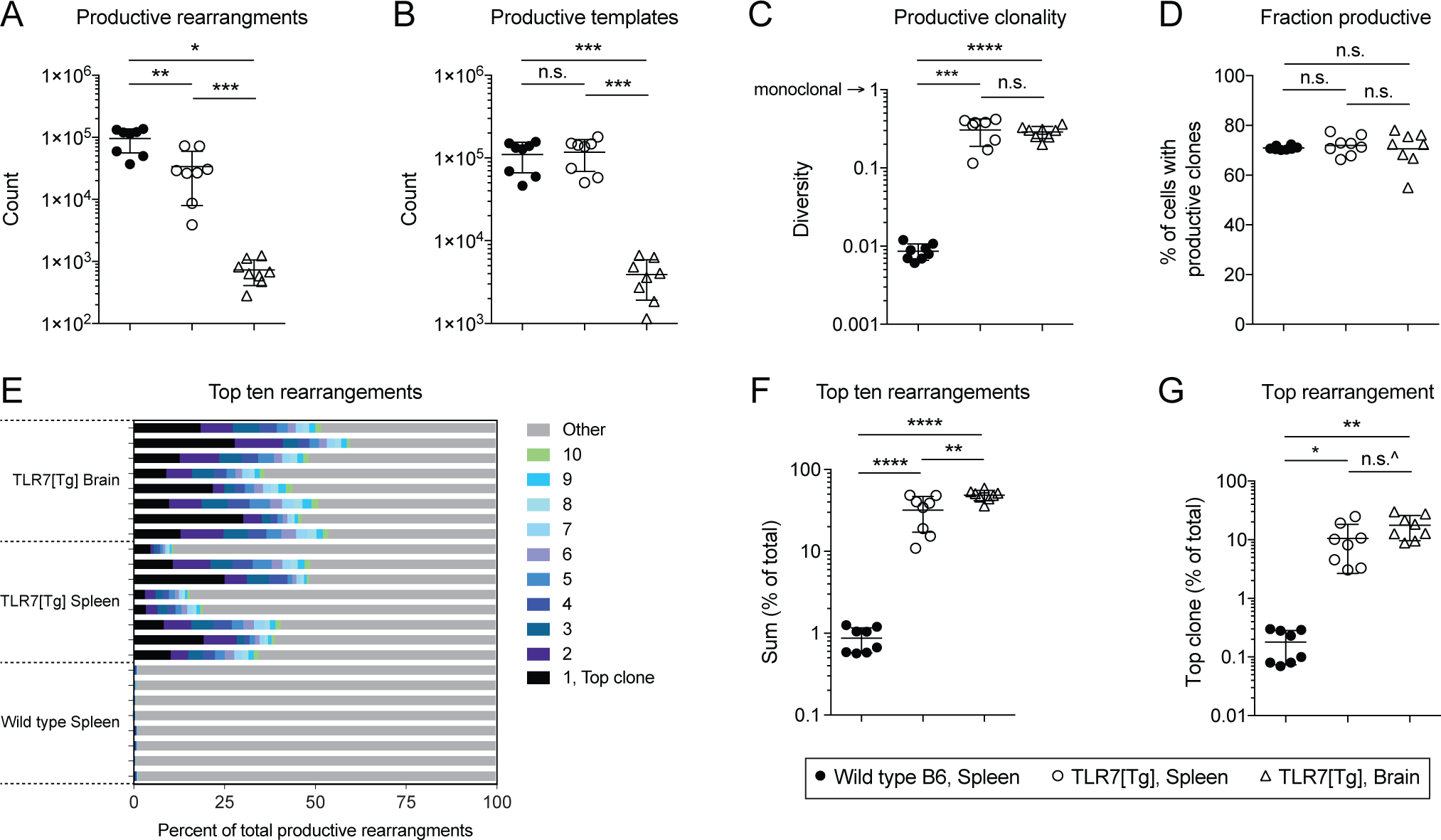
CD8+ T cells from lupus-prone mice demonstrate substantial oligoclonality. (**A-D**) Overview of total TCRβ rearrangements from wild type B6 and TLR7[Tg] CD8+ T cells from Figure 1. (**A**) The count of unique rearrangements in each sample that are in-frame and do not contain a stop codon. (**B**) The sum of templates for all productive rearrangements in the sample. (**C**) Clonality measure for each sample calculated over all productive rearrangements. (**D**) The fraction of productive templates among all templates. (**E-G**) The frequency (**E**) and sum (**F**) of the top ten clones and the top clone (**G**) for each sample relative to all rearrangements. Error bars inducate mean + s.d. ^P = 0.08, *P ≤ 0.05, **P ≤ 0.01, ***P ≤ 0.001, and ***P ≤ 0.0001, student’s t test.

An analysis of the most abundant ten clones in the wild type control reveals an average summative frequency of less than 1%, indicative of a polyclonal response (Fig. 2E, 2F). By comparison the strong oligoclonality of both splenic and brain CD8^+^ T cells from TLR7tg mice can also be seen in the summative frequency of the top ten clones in each animal, which can amount to 60.5% of the repertoire (Fig. 2E, 2F). In this case, the sum of the top clones in the TLR7tg brain with respect to the entire repertoire was statistically higher relative to the sum of the top clones in the TLR7tg spleen (Fig. 2F). Remarkably, the top unique rearrangement in the total TLR7tg CD8^+^ T cell pool was 30.1% of all productive templates, while this number never exceeded 0.4% for the wild type animals (Fig. 2G).

### Skewed CDR3 length and VDJβ gene usage in TLR7tg CD8^+^ T cells compared to wild type

An established method of assessing the relative clonality in a given lymphocyte repertoire is through an analysis of the various CDR3 lengths within the population. A Gaussian distribution of the CDR3 lengths is indicative of polyclonal responses (21). Consistent with this we found that CDR3 sequence lengths of CD8^+^ T cells from healthy control mice demonstrated a very strong fit to a Gaussian curve centered on fourteen amino acids (Fig. 3A and 3B, *top*), which is consistent with a recent comprehensive analysis including B6 control animals (22). By comparison the CDR3s from TLR7tg splenic CD8^+^ T cells had a substantial divergence from Gaussian distribution (Fig. 3A and 3B, *middle*). Additional variance was found in brain isolated cells (Fig. 3A and 3B, *bottom*). Importantly, despite a narrowing repertoire, the CDR3 sequence distribution across all TLR7tg samples tested is not uniformly directed towards a specific set of rearrangement processes; each of the eight lupus-prone animals exhibit a uniquely skewed abundance of CDR3 lengths. There is, however, a similar pattern of CDR3 length usage between shared spleen and brain samples within each of the same TLR7tg animals (Fig. 3A and 3B, *middle* and *bottom*, sequentially matched bars *left* to *right*).

**FIGURE 3.**
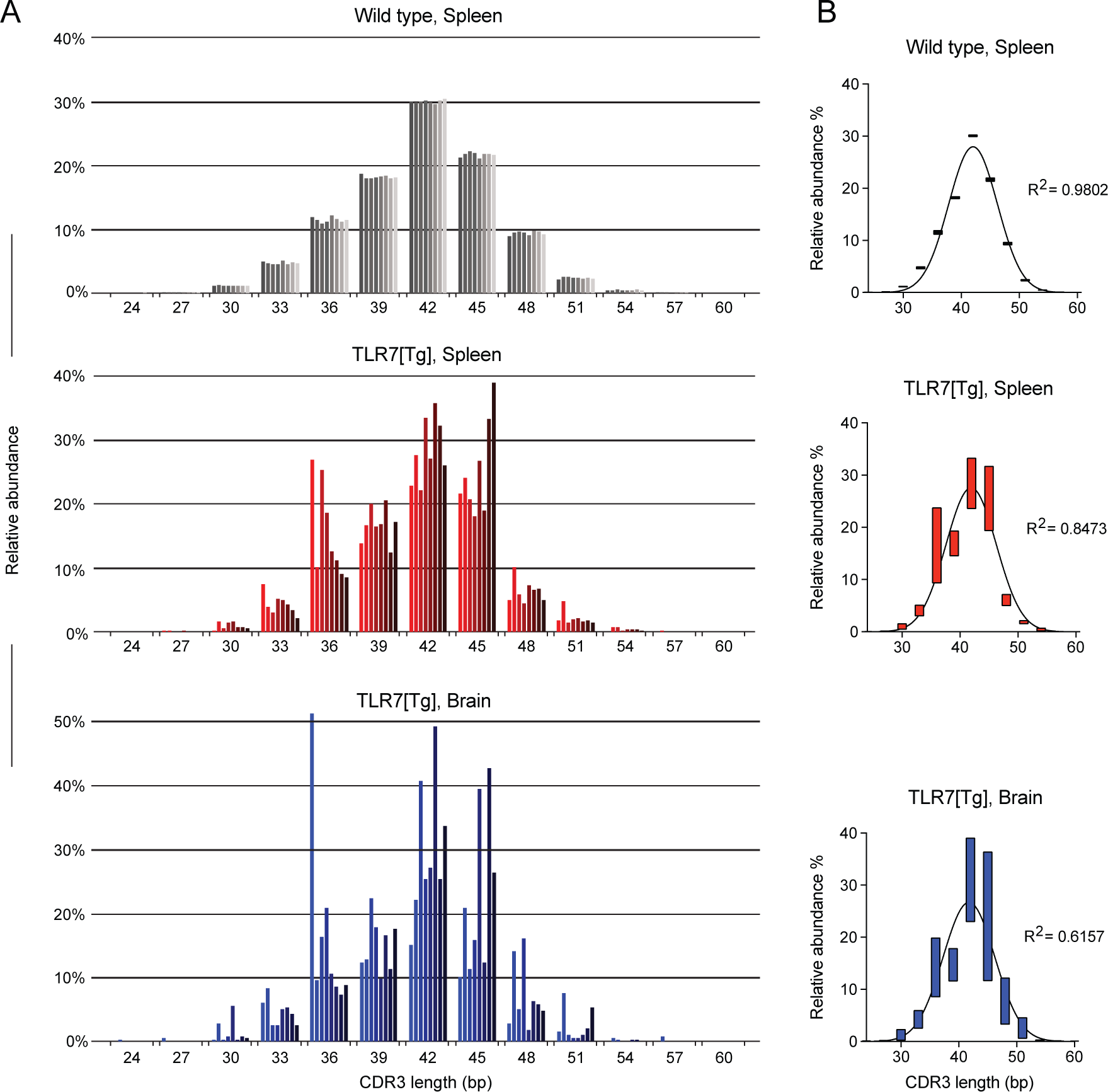
Variable CDR3 legnths support the notion of oligoclonality in CD8+ T cells of lupus-prone mice. (**A**) Relative abundance and distribution of CDR3 nucleotide sequence lengths (bp) from purified CD8+ T cells of wild type B6 spleen (shades of grey), TLR7[Tg] spleen (shades of red), and TLR7[Tg] brain (shades of blue). Each bar represents a separate sample of the indicated genotype and tissue. For each possible CDR3 length the order of the eight bars from left to right is matched for spleen and brain samples from TLR7[Tg] mice. (**B**) Average distribution analysis of compiled samples (n=8) from (A). Bars indicate minimum and maximum values centered on the mean. R^2^ values indicate the fit of each curve to normal Gaussian distribution.

We next analyzed VDJβ gene usage in CD8^+^ T cells from wild type and TLR7tg mice. The results in Figure 4 include total Vβ gene segment usage and are presented with two forms of nomenclature. The international Im<u>M</u>uno<u>G</u>ene<u>T</u>ics information system (IMGT) is the current standard used since 2000 and includes all pseudogenes in the Vβ gene region (23), though many publications and commercial antibody suppliers still adhere to the original distinctions (24). Naïve animals rely on a relatively small pool of TCRβ genes in the makeup of their peripheral repertoire (25). Our analysis of the VDJβ gene makeup of the CD8^+^ T cells in wild type B6 mice was consistent with these published results, with the majority of cells utilizing Vβ12 and Vβ13 (Fig. 4A), Dβ1 (Fig. 4B), and Jβ2.7 (Fig. 4C) gene segments. In contrast, cells from the spleen of TLR7tg mice showed highly variable VDJβ usage (Fig. 4A-C), and significant divergence from common gene segments such as Vβ12, and Jβ 2.5 (Fig. 4D). The brain CD8+ T cells from TLR7tg mice also used statistically less Vβ12, and Jβ 2.5. In place of these common Vβ and Jβ genes both the spleen and brain cells from lupus-prone mice did use other genes, though no consistent pattern emerged across those samples. In fact, substantial mouse-to-mouse variability of VDJβ genes used was evident in both TLR7tg splenic and brain CD8^+^ T cell populations (Fig. 4D).

**FIGURE 4.**
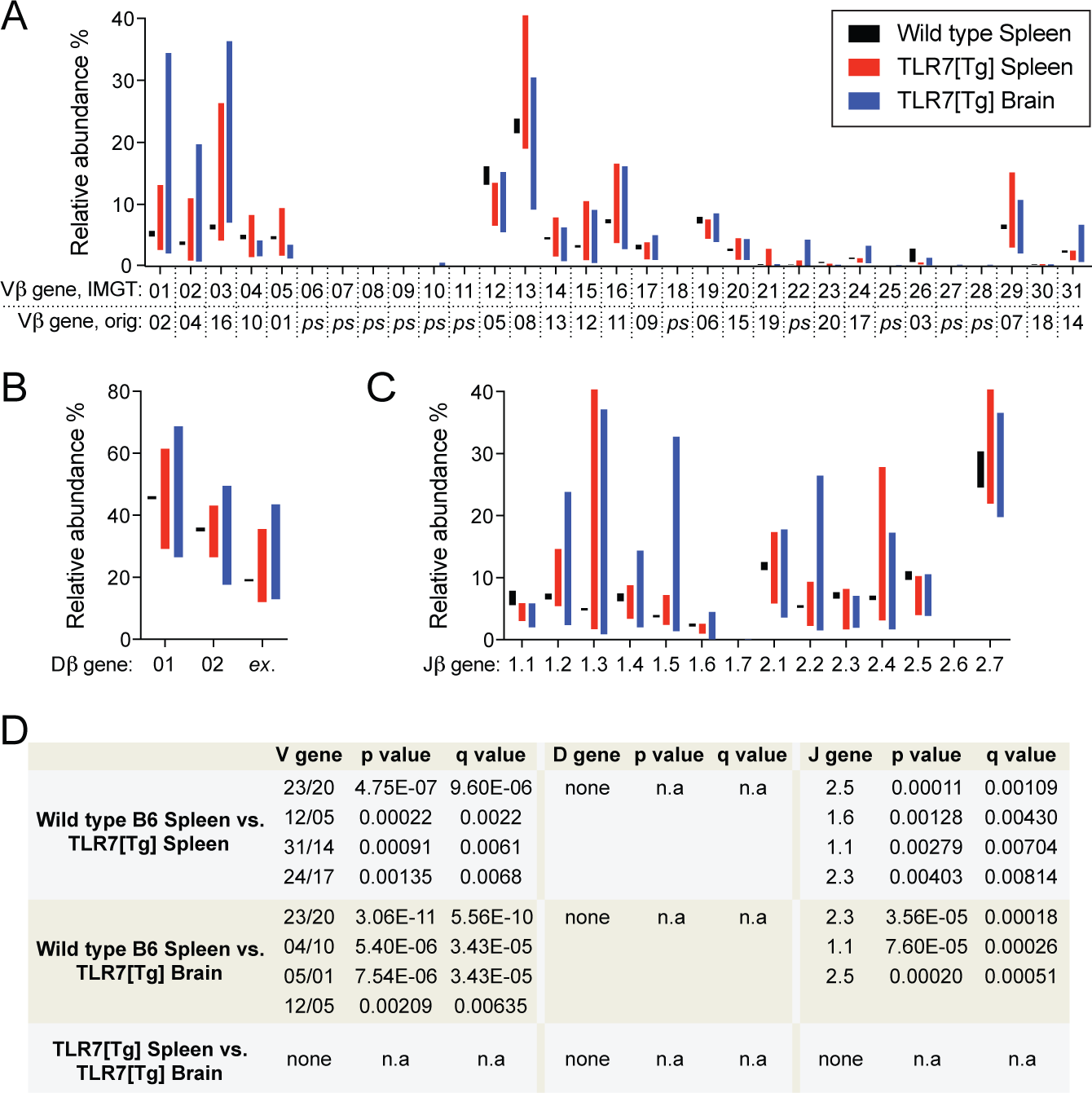
Highly variable VDJβ gene usage in lupus-prone CD8+ T cells. (**A-C**) Vβ gene (**A**) usage for CD8+ T cells from wild type B6 spleen, and TLR7[Tg] spleen and brain. Equivalent IMGT and original (orig.) nomenclature for Vβ genes included. *ps*, pseudogene. (**B**) Dβ gene and (**C**) Jβ gene usage shown as in (A). Bars indicate minimum and maximum values centered on the mean. (**D**) Statistical analysis of discovered significant differencences between indicated groups for (A-C). Multiple t test grouped analysis used to generate p values and false discovery rate-adjusted q values. These results are presented with two forms of nomenclature: A newer and comprehensive one including pseudogenes from the international ImMunoGeneTics information system (IMGT), and the original nomenclature still common.

### The peripheral and brain-infiltrating CD8^+^ T cell pools within the same lupus-prone mouse show substantial CDR3 nucleotide sequence sharing

To better understand the substantial mouse-to-mouse variations in TLR7tg samples despite similar levels of oligoclonality we analyzed absolute CDR3 nucleotide homology across all in-frame rearrangements from every animal tested between spleen and brain tissues of both genotypes (Fig. 5A-F). We identified a small number of shared identical CDR3 sequences between various wild type B6 splenic samples (Fig. 5A, 5E, 5F). Unexpectedly, a significant reduction in sharing across splenic TLR7tg mice was evident (Fig. 5B, 5E, 5F). We found additional reduction in such sharing across all brain isolated populations (Fig. 5C, 5E, 5F), in some cases down to only a single shared sequence between two genetically identical animals. Within the same animal we did find considerable instances of complete nucleotide homology in the CD8^+^ T cells between spleen and brain tissues (Fig. 5D, 5E), including sharing of the most abundant clone between seven of the eight TLR7tg mice (Fig. 6A). These results indicate that lymphocyte expansion in a chronic-inflammatory autoimmune setting can give rise to an oligoclonal population of CD8^+^ T cells in which the same clones are likely to be found in both lymphoid and non-lymphoid tissues, but only within the same animal.

**FIGURE 5.**
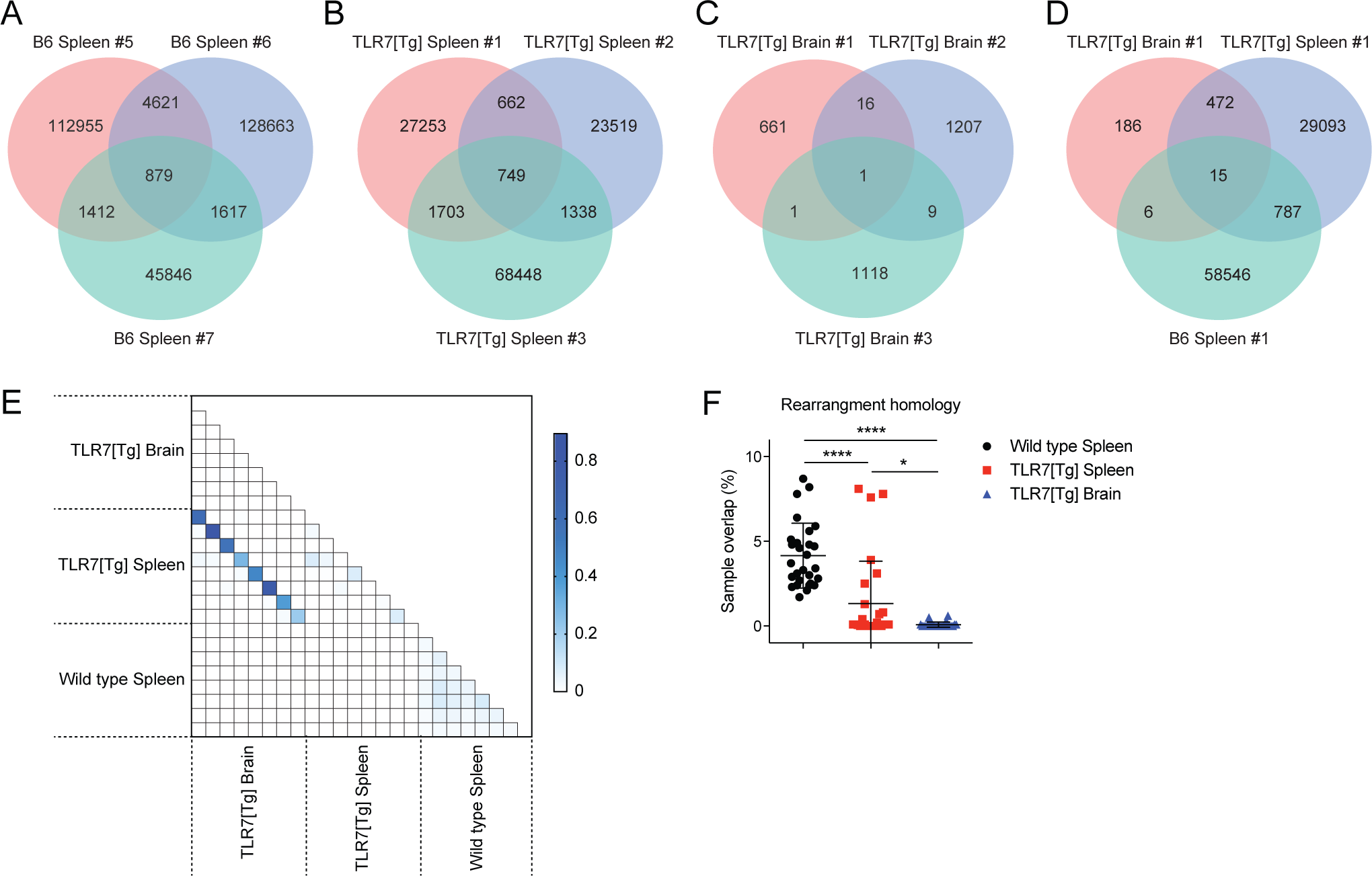
The CD8+ T cell repetoire is largely similar between the spleen and brain of the same lupus-prone animal. (**A-D**) Venn-diagram analysis of CDR3 nucleotide sequence homology from three wild type B6 spleens (**A**), three TLR7[Tg] spleens (**B**), three TLR7[Tg] brains (**C**), and across all three groups (**D**) comparing in-frame reads. (**E**-**F**) CDR3 nucleotide sequence overlap across all in-frame reads of samples (n = 8) from indicated groups calculated by Morisita Index (**E**) and summarized for each group (**F**). Error bars inducate mean + s.d. *P ≤ 0.05, **P ≤ 0.01, ***P ≤ 0.001, and ***P ≤ 0.0001, student’s t test.

**FIGURE 6.**
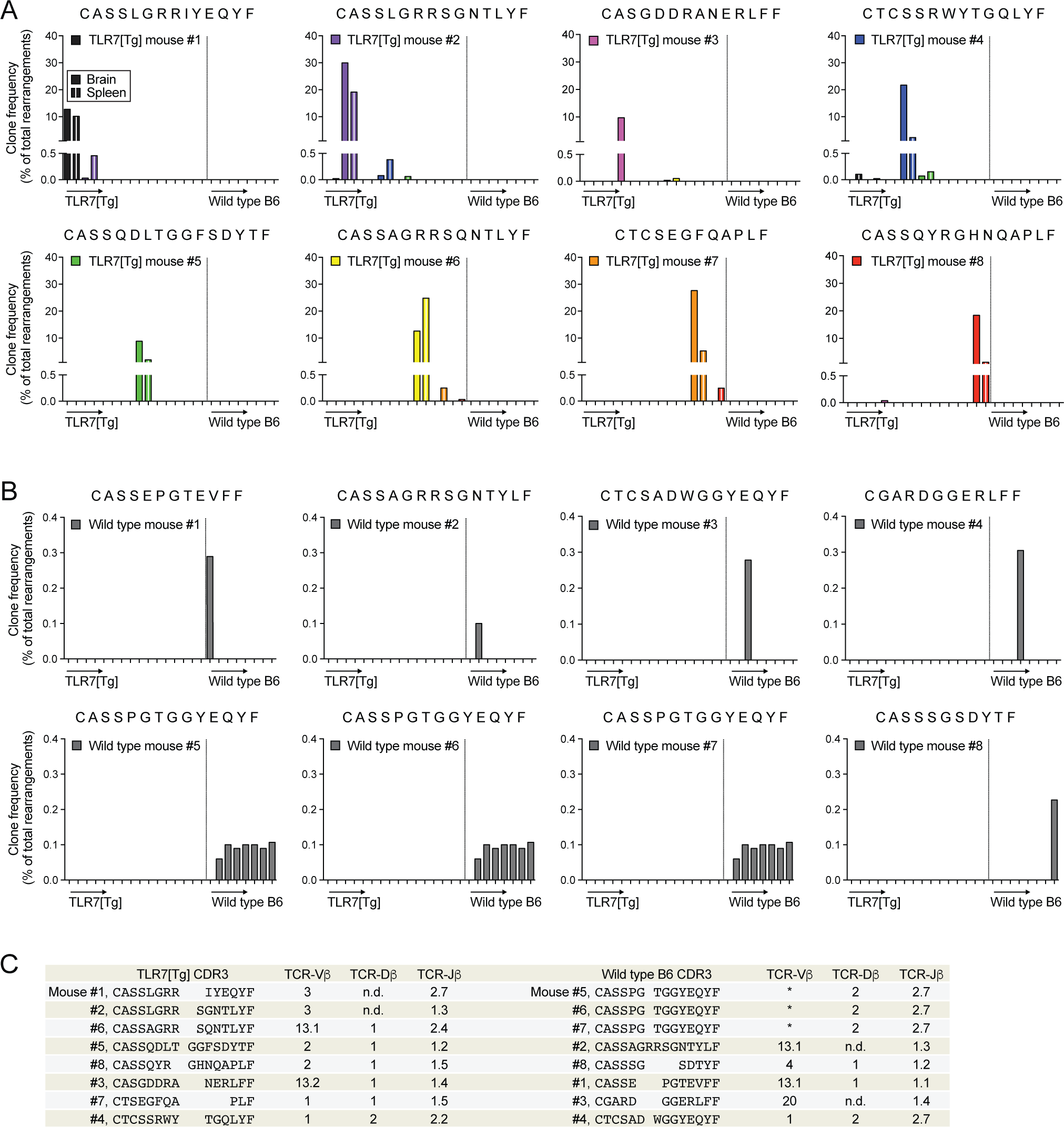
Regions of sequence homology across dominant clones in TLR7[Tg] animals. (**A**, **B**) The CDR3 sequence of the dominant CD8+ T cell clone isolated from the brain of each TLR7[Tg], n = 8 (**A**) or spleen of each wild type B6, n = 8 (**B**) mouse is shown. The frequency at which each of these top clones appears across all animals is presented. (**C**) CDR3 alignment along with V, D, and J gene usage for each top clone in TLR7[Tg] or wild type B6 mice. An asterisk (*) indicates an instance in which multiple nucleotide sequences encoding different Vβ genes generated identical CDR3 amino acid sequences.

### Dominant clones across lupus-prone mice share regions of sequence homology, but are distinct the most common clones in healthy control animals

Remarkably, while the most abundant single clone in each wild type control sample was only between 0.1% to 0.4% of the entire population, in TLR7tg mice this number reached between 10% and 30% for splenic samples (Fig. 6A). Similar percentages were found for brain-isolated CD8^+^ T cells. The CDR3 amino acid sequences of the top rearrangements from wild type and TLR7tg mice were largely distinct (Fig. 6B), indicating that the CD8^+^ T cells selected by the environment in lupus-prone mice are different from the top clones in wild type controls.

Despite the strong oligoclonality we witnessed in the lupus-prone samples, complete nucleotide homology across various TLR7tg mice was rare (Fig. 5). A possible scenario is that similar but not identical CDR3 sequences might be selected by the same antigen within expanded T cell populations in these animals. We identified several highly similar dominant clones, particularly those from TLR7tg mouse #1, #2, and #6, which contained up to 93% amino acid sequence homology while representing nearly one-third of the CD8 repertoire in each animal (Fig. 6A, 6C). Interestingly, even though there was such a high degree of sequence homology in these clones, their VDJ gene usage was largely distinct. These results collectively indicate that clonal expansion in TLR7tg lupus-prone mice selects for particular dominant rearrangements that share regions of sequence homology despite not deriving from the same mature parent thymocyte.

## Discussion

The specificities and clonality of the lymphocytes thought to be involved in systemic lupus are not a well understood aspect of the disease, particularly as they concern CD8^+^ T cells. Much of the data in the current study supports the notion of oligoclonality in the TLR7tg CD8^+^ T cells compared to wild type controls; a narrowing of the total number of unique clones (Fig. 2), a non-Gaussian distribution of CDR3 length abundance (Fig. 3), and skewed VDJβ gene usage (Fig. 4). The level of clonality we observe in the TLR7tg animals, with a single clone composing up to 30% of the entire T cell pool, is similar to what has been published in CD8-dependent models of viral infection such as LCMV (26) and influenza (27). An important distinction between the TLR7tg lupus-prone mouse model and viral infection systems is the scope of available antigens to which the T cells can respond. As a result, only three predominant CD8^+^ T cell clones have been described against the Armstrong strain of LCMV (26). Importantly, while each of these clones contain unique VDJ genes, CDR3 lengths, and nucleotide sequences, they are consistently present across all infected animals. In our lupus model system, despite a similar degree of oligoclonality in the CD8^+^ T cell pool there is an unexpectedly broad and variable usage of VDJ genes, CDR3 lengths, and sequences. The importance of this distinction remains unclear, though it is likely indicative of a heterogeneous response against a group of self-antigens.

In a previous study, we showed that the CD8^+^ T cells in TLR7tg mice are activated effector cells that, compared to other lymphocytes in the animal, could traffic to and take up residence in the brain (5). Activated lymphocytes gain adhesive properties allowing them to attach to blood vessel endothelium of tissues into which they will migrate (28).

The most highly activated CD8^+^ T cells in the TLR7tg mice upregulated many adhesion and trafficking markers (5), and so we posited that these cells would enter the brain and expand in response to some antigen specific signals. Unexpectedly, our current data identified a markedly similar TCRβ repertoire between the peripheral and brain-resident CD8^+^ T cell pools in each respective animal (Fig. 5D, 5E, 6A). It is clear that of all the possible comparisons we could make across tissues and genotypes within a single animal, the strongest degree of sequence homology existed between cells within different tissues of the same animal. We conclude from these results that the pool of expanded, oligoclonal, mature lymphocytes we identified in both the spleen and brain of TLR7tg mice are develop in response to the same selective pressures, though it remains unclear if the origin of such selection is from peripheral or brain antigens.

Nucleotide homology of the antigen-determining CDR3 region can provide some information about the CD8^+^ T cell repertoire. While absolute homology across TLR7tg mice is rare, particularly in the brain, it is possible that the high degree of similarity identified across some of the top rearrangements is indicative of different clones arising with similar specificities, perhaps for some common self-antigen. This is possible even in cases when the different VDJ genes are used to make two unique rearrangements. For example, while a close relationship between the CDR3 specificity and the structure of the antigen receptor exists, and most of the T cells with the same MHC:peptide specificity were found to use the same αβ-chain combinations (29, 30), examples exist of various insulin-specific T cell clones and hybridomas using different Vα and Vβ genes (31, 32).

One interesting implication of our data relates to the notion that shrinking receptor diversity resulting from clonal expansion in individuals with chronic disease such as systemic lupus could promote what has been referred to as an ‘immune risk phenotype’ (33) during senescence. Such a phenotype is predictive of mortality in aged individuals who are left more susceptible to constant immune insult from foreign pathogens. The narrowing of the lymphocyte repertoire, which is thought to impact CD8^+^ T cells more than CD4^+^ T cells, decreases immune defense particularly during senescence in both mice and humans (33-35). Because we see a narrowing lymphocyte repertoire in the context of autoimmune disease, it will be important in the future to use our lupus-prone mice and other such models to study the impact of chronic systemic disease on the ability of the organism to mount both effective primary and secondary immune responses to pathogens.

The antigen specificity of the CD8^+^ T cells we identified remains unknown, making it hard to speculate what might be driving the mouse-to-mouse differences despite similar overall clonality. Certainly, the frequency and sequence homology of the most abundant clones suggests a strong selective pressure indicative of some self-antigen-driven expansion of CD8^+^ T cells. However, how the top clones of the CD8^+^ T cell pool end up as part of the response in lupus-prone mice to begin with and why the repertoire is so variable between animals remains unclear and warrant further investigation.

## Acknowledgements

We thank Bethany Scott for technical assistance with animal care and vascular labeling. We are grateful to Kevin Holmes and the NIAID RTB Cytometry section for assistance isolating lymphocytes for sequencing. This research was supported by the Intramural Research Program of the NIH, NIAID.

